# Environmental DNA metabarcoding as an effective and rapid tool for fish monitoring in canals

**DOI:** 10.1101/498451

**Authors:** A. D. McDevitt, N. G. Sales, S. S. Browett, A. O. Sparnenn, S. Mariani, O. S. Wangensteen, I. Coscia, C. Benvenuto

## Abstract

Environmental DNA (eDNA) metabarcoding has revolutionized biomonitoring of aquatic habitats. Man-made canal systems are among the least-studied environments in terms of biodiversity in Britain. Here we focus on a case study along an English canal comparing eDNA metabarcoding with two types of electrofishing techniques (wade-and-reach and boom-boat). In addition to corroborating data obtained by electrofishing, eDNA provided a wider snapshot of fish assemblages. Given the semi-lotic nature of canals, we encourage the use of eDNA as a fast and cost-effective tool to detect and monitor whole fish communities.

---

England’s biodiversity depends on diverse habitats that are currently protected as SSSI (Sites of Special Scientific Interest). Among these designated areas, there are several canal systems, which are monitored for habitat quality and the occurrence of certain indicator species (Mainstone *et al.*, 2018). However, despite there being over 3,000km of canals in the United Kingdom, little has been done to assess their entire biodiversity (Natural England, 2011). Improvement of open freshwater habitats is considered challenging, and long-term routine monitoring of canal systems is critically important, as these can be key in the assessment of invasive, migratory and/or endangered species, as well as safeguarding against the spread of diseases through early detection of pathogens.

Traditionally, teleost populations have been monitored through live capture and subsequent morphological identification of specimens (Hill *et al.*, 2005). However, these practices are intrusive, can compromise the health of targeted species and induce stress (Goldberg *et al.*, 2016). The selectivity of equipment used during traditional surveying practices can also lead to inaccuracies when monitoring freshwater ecosystems because specialized equipment can exclude the sampling of specific species (due to size, microhabitat use, and low abundances), thus leading to an insufficient representation of the community (Evans & Lamberti, 2017). Furthermore, the limited access to specialized equipment (such as electrofishing gear) and funding can make traditional surveys expensive and restrictive (Shaw *et al.*, 2016).

Environmental DNA (eDNA) metabarcoding has emerged as an innovative and effective biodiversity monitoring tool that enables the rapid classification of multiple taxa without the assistance of a taxonomist or local fishing knowledge (Taberlet *et al.*, 2012). A cost-effective, fast and non-invasive eDNA protocol could prove extremely useful to provide a constantly updated and broad monitoring of the aquatic biodiversity of canals and its changes through time. The present study focuses on the detection capability of eDNA metabarcoding compared to two different types of electrofishing for the detection of fish species along a stretch of the Huddersfield Narrow Canal in the UK (Fig. 1A). This canal is a designated SSSI and has very limited data available in terms of fish assemblage and biodiversity.

**Figure 1.**
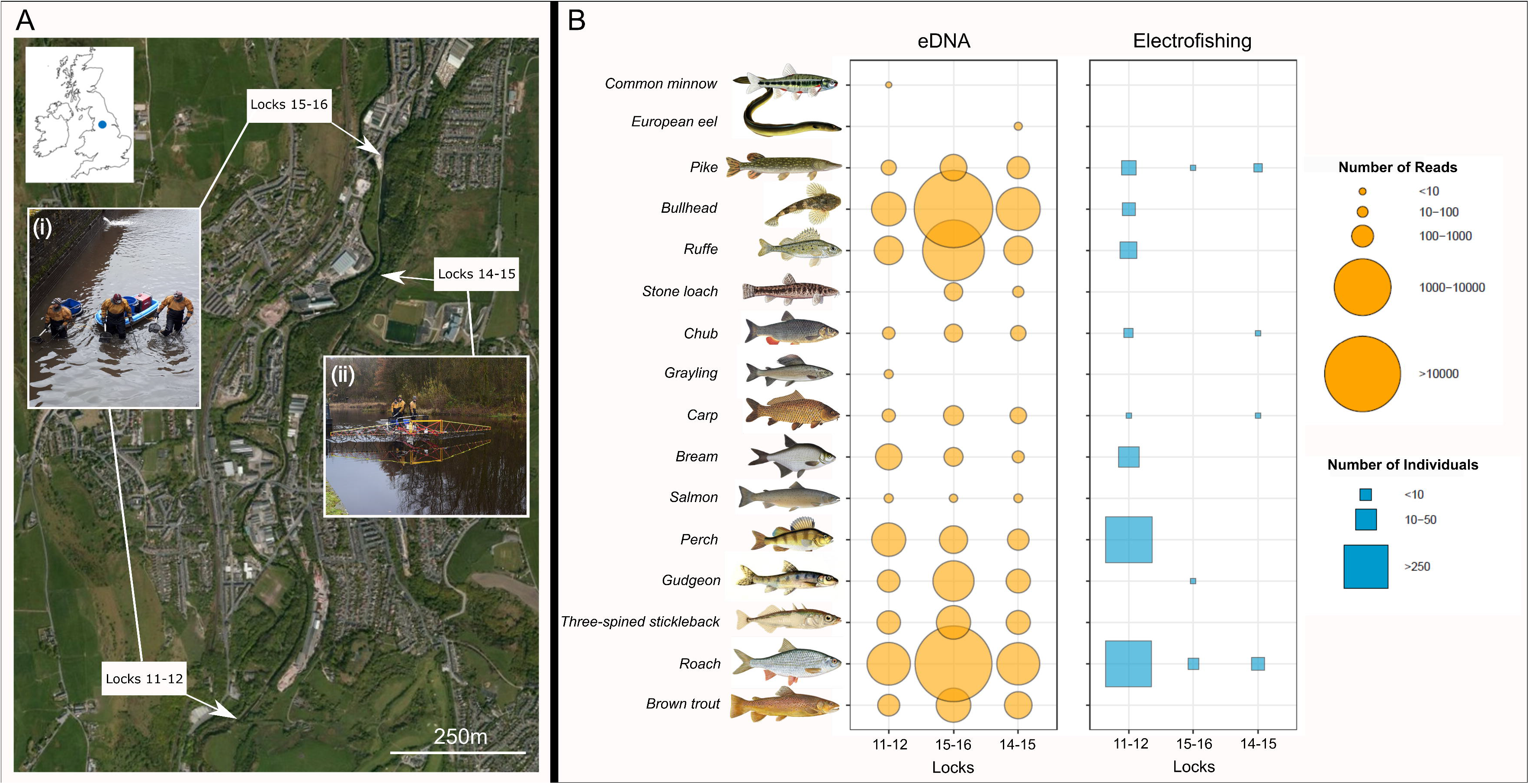
Map of the study area showing sampling locations for electrofishing (wade-and-reach (i) and boom-boat (ii)) and eDNA between Locks 11-16 of the Huddersfield Narrow Canal (A). A bubble graph (B) is used to represent presence-absence and categorical values of the number of reads retained (after bioinformatic filtering) for eDNA (water and sediment combined) and the number of individuals caught for electrofishing for 16 fish species. Fish illustrations not shown to scale.

Three stretches of the canal were chosen for a Canal and River Trust-commissioned fish survey in December 2017/January 2018. Electrofishing surveys were conducted using two methods: ‘backpack’ electrofishing with water levels lowered and surveyors wading through the canal bed between Locks 11-12 and 15-16 (Fig. 1A(i)) and a boom boat with water levels maintained between Locks 14-15 (Fig. 1A(ii)). Three sweeps were undertaken at each stretch and fish were identified to species level. One to 16 hours prior to these surveys being conducted (and before water levels were lowered), water temperature and pH were measured, and water (5 × 2L) and sediment (3 × ~10g) samples were taken from each of the three stretches of the canal. Water samples were filtered (250-400ml) within three hours of sampling in a decontaminated laboratory using Sterivex 0.45μM filters that were then kept at -20°C; sediment samples were stored in 100% ethanol at room temperature. To avoid cross contamination between samples, appropriate decontamination measures/precautions were taken: gloves were worn at all times, equipment and surfaces were treated with bleach (10%) and three field blanks were also analysed.

DNA was extracted from the water samples using the DNeasy PowerWater Kit and from the sediment samples using the DNeasy PowerMax Soil Kit (both Qiagen) in the lab. All field blanks were extracted first, and extractions were completed following the manufacturer’s protocol. Due to the nature of the sediment it was not always possible to collect 10g free of macroremains. Amplification of a fragment of the mitochondrial 12S rRNA gene was conducted using the MiFish 12S primer set (Miya *et al.*, 2015) and library preparation were conducted according to the protocol described in Sales *et al.* (2018). A total of 29 samples (including collection blanks and laboratory negative controls) were sequenced in a single multiplexed Illumina MiSeq run along with samples from a non-related project (not included in this study). See the Supplementary Material for details on the bioinformatic analyses.

A total of nine species were identified with the two electrofishing methods. With the boom boat, pike (*Esox lucius*), roach (*Rutilus rutilus*), chub (*Squalius cephalus*) and carp (*Cyprinus carpio*) were captured between Locks 14-15. Using the other electrofishing method (wade-and-reach) between Locks 11-12 and 14-16, perch (*Perca fluviatilis*), gudgeon (*Gobio gobio*), bream (*Abramis brama*), ruffe (*Gymnocephalus cernuus*) and bullhead (*Cottus gobio*) were captured in addition to the previous four species. Only roach and pike were captured across all three electrofishing sessions (Fig. 1B).

A total of 104,055 sequence reads (after all filtering steps) were retrieved, allowing for the detection of 16 species in the eDNA survey. All nine species from the electrofishing survey were identified, with the addition of brown trout (*Salmo trutta*), common minnow (*Phoxinus phoxinus*), European eel (*Anguilla anguilla*), grayling (*Thymallus thymallus*), salmon (*Salmo salar*), stone loach (*Barbatula barbatula*) and the three-spined stickleback (*Gasterosteus aculeatus).* The results provided by eDNA were more consistent, with 12 out of the 16 species being detected in all three sampling sessions (Fig. 1B). Electrofishing failed to detect seven species, and the selectivity of the method may hamper the detection of species difficult to capture due to their morphological or behavioural characteristics (small body size fish species such as *P. phoxinus, G. aculeatus,* or solitary and nocturnal fish such as *B. barbatula*).

Due to the expected relatively fast degradation of DNA molecules (Seymour *et al.* 2018), the detection of species through this method suggests their recent presence and provides an overview of the contemporary fish community. However, eDNA molecules might persist in the water column for more than a few days and thus, allow the detection of transient species not necessarily present in the system at the collection time (Dejean *et al.*, 2011, Thomsen *et al.*, 2012). DNA molecules can be transported long distances (Barnes & Turner, 2016) so fish may be detected some distance from their occurrence (Jane *et al.*, 2015) or even originating from different sources. Therefore, the detection of the rare species (e.g. eel and salmon) in this study could be due to an external source. Putative false positives should be taken into account and carefully analysed before drawing a conclusion about the occurrence of these species in the Huddersfield Canal, and to understand their origin (e.g. endogenous or exogenous, regional or local).

As previously shown by Shaw *et al.* (2016), eDNA obtained from the water column yielded better results when compared to sediment samples with 14 out of 16 species recovered, and sediment samples outperformed water samples only by detecting eel and minnow. Environmental DNA recovered from sediment samples allowed the detection of only five species (eel, brown trout, salmon, minnow and stone loach; Table S1). The type of sediment has an impact on how much eDNA is retained (Shogren *et al.*, 2016), meaning that the compacted nature of the sediment in this canal could lead to low eDNA retention.

While many studies have shown the advantages of using eDNA metabarcoding in lotic (flowing streams and rivers; Balasingham *et al.*, 2018) and lentic (still lakes and ponds; Harper *et al.*, 2018; Hänfling *et al.*, 2016; Perez *et al.*, 2017) systems, they also raise concerns about the influence of flow in DNA dispersal in fast running water and the need to sample multiple locations in lentic waters. Canals represent man-made environments with a semi-lotic regime and regulated flow, which minimize the risk of detection of species present too far away, while at the same time allowing enough water movement to reduce the need of extra sampling akin to that undertaken in lentic systems. Here we showed that environmental DNA corroborates the data obtained by electrofishing, but also provides a wider snapshot of fish assemblages (Pont *et al.*, 2018). While traditional methods cannot be replaced when investigating size, age class distribution, and, for now, abundance, we find that the power, speed and cost-effectiveness of eDNA metabarcoding may often represent a highly efficient tool to assess and monitor whole fish communities in canal systems.

## Acknowledgements

We thank The Peoples Postcode Lottery and the University of Salford for financial support. We are grateful to Thomas King and Linda Butterworth at the Canal and River Trust for advice and discussions, and to MEM Fisheries for access to their data from the electrofishing surveys.

